## Supplementary figures and images for "Comparative gene annotation and orthology assignments across 301 species of Drosophilidae"

### S1 Fig

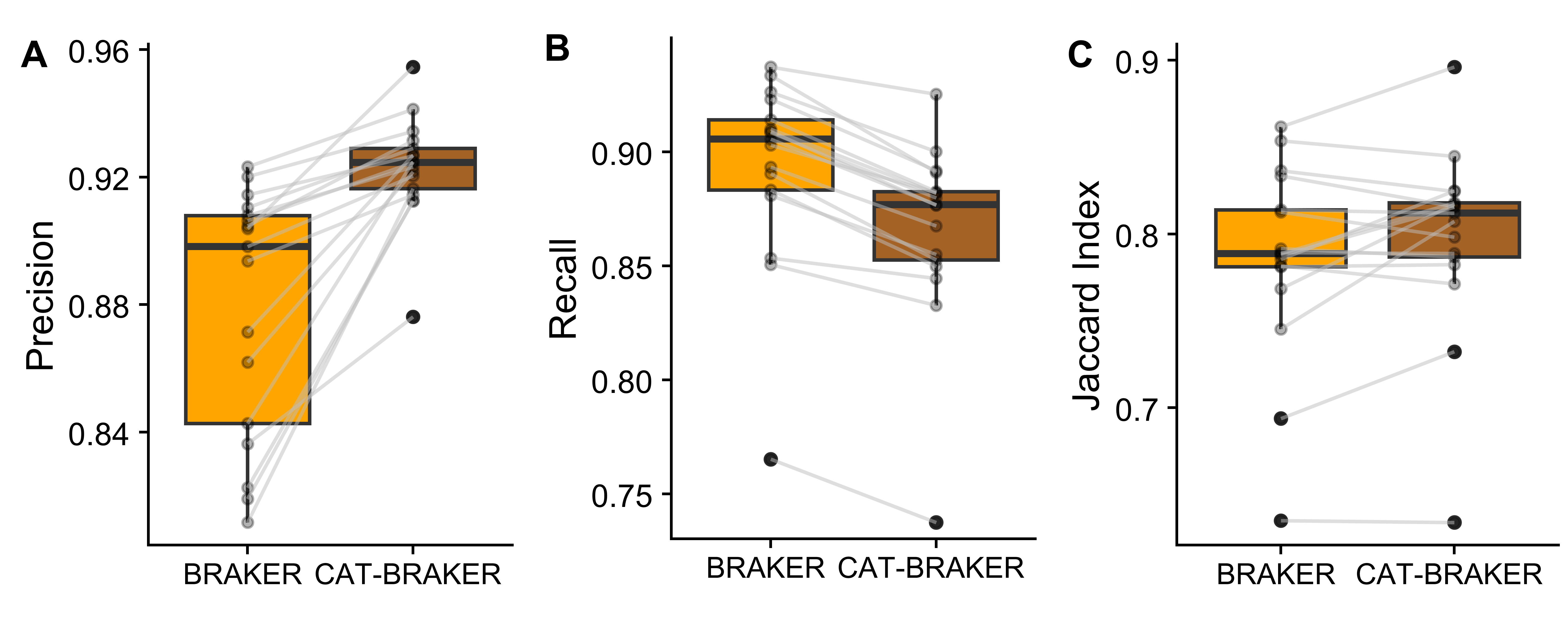

### S2 Fig

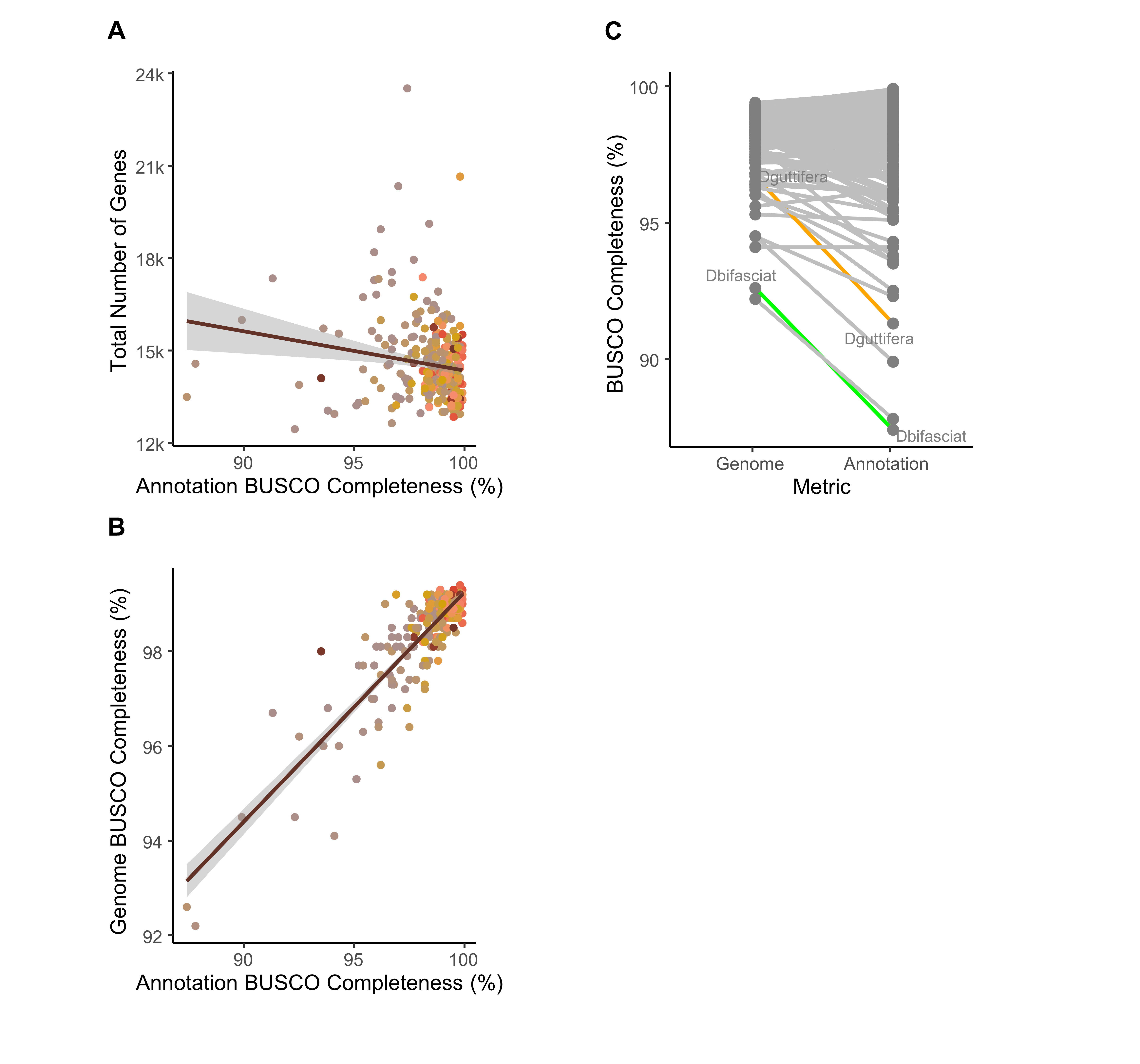

### S2 File

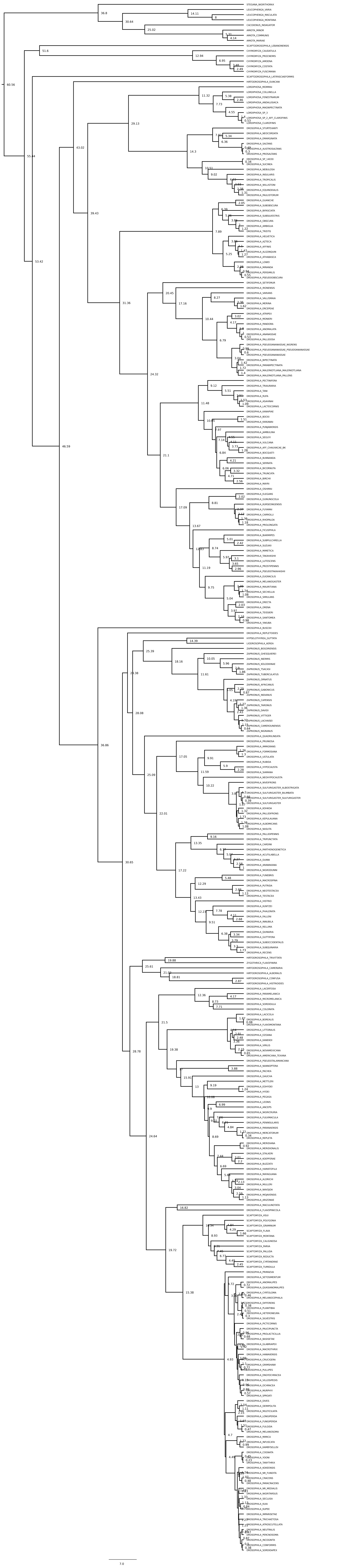

### S3 Fig

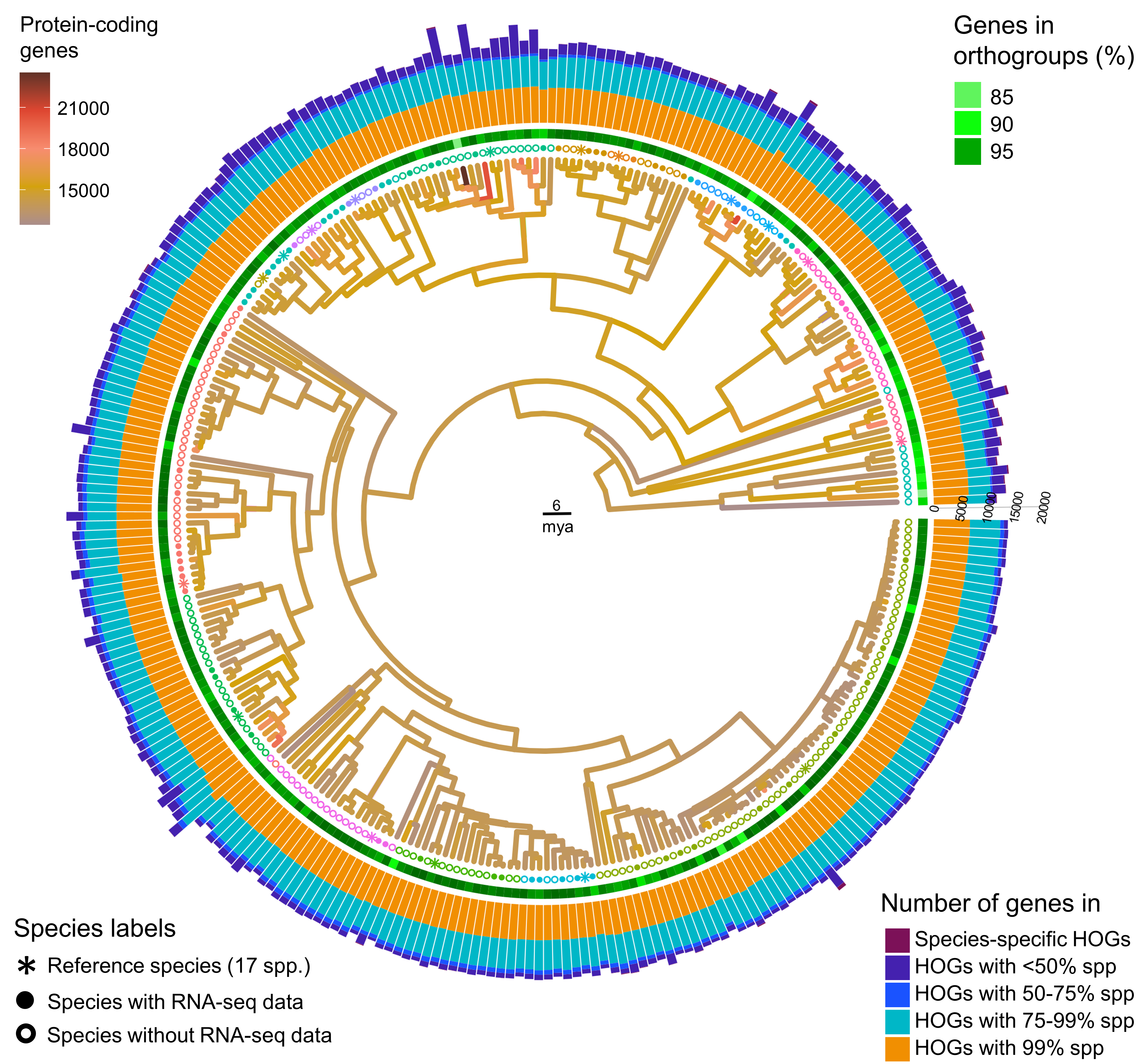

### S3 File

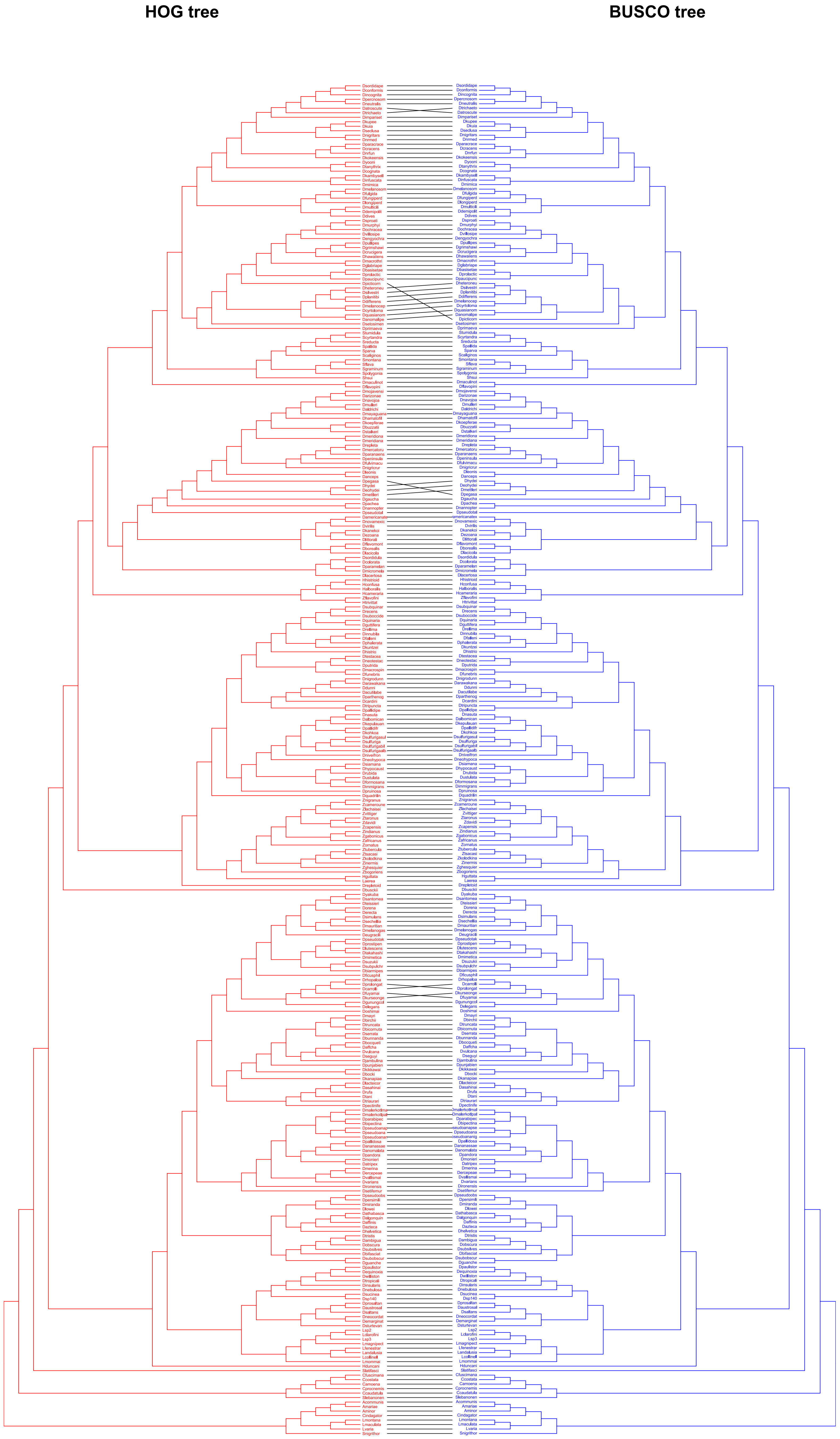

### S4 Fig

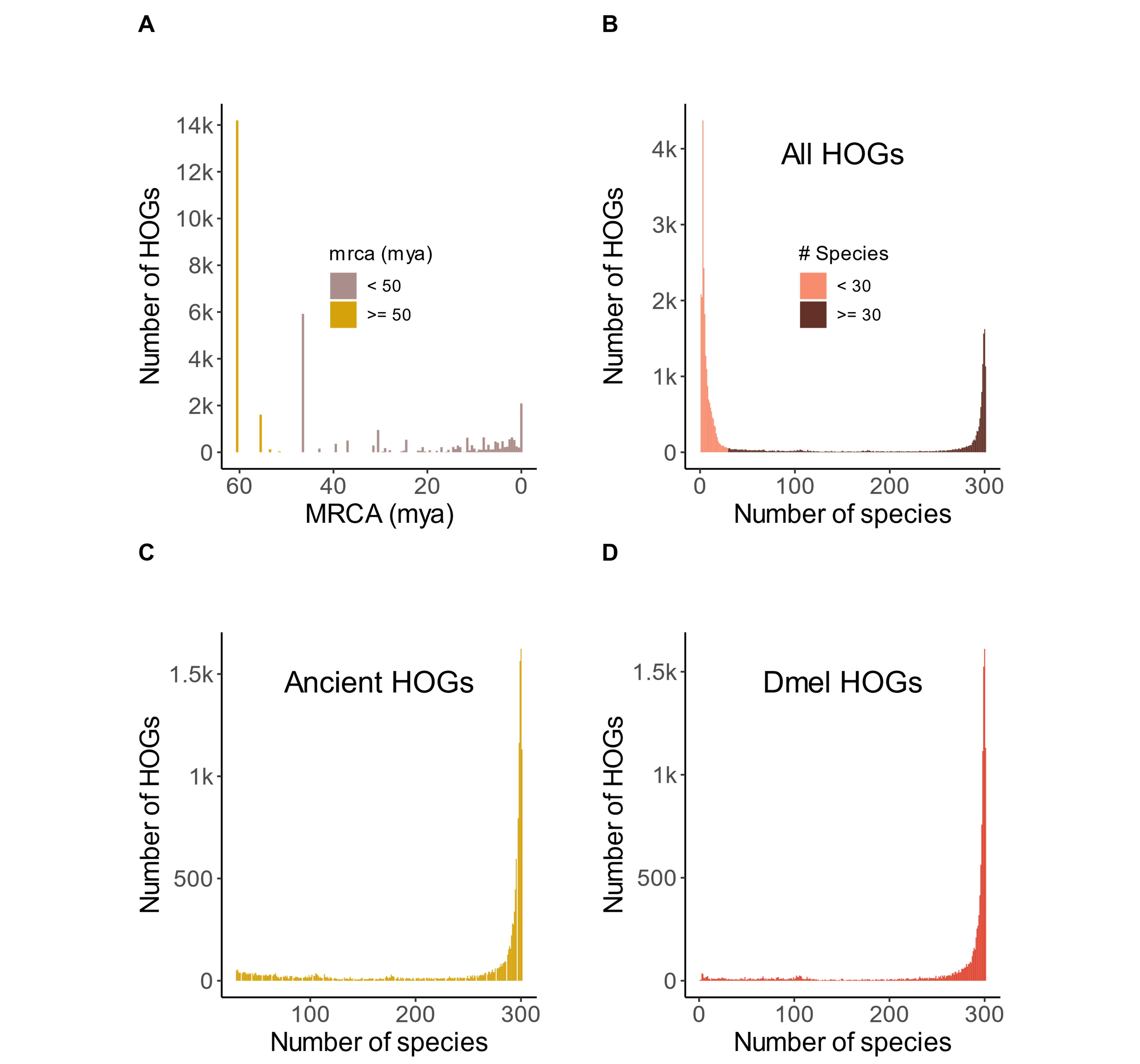

### S4 File

#### HOG tree

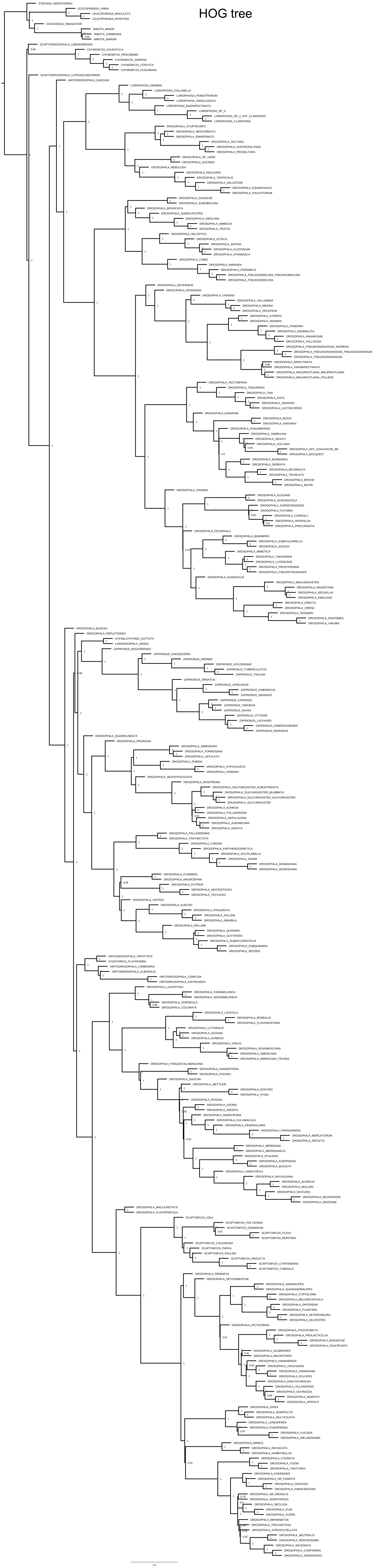

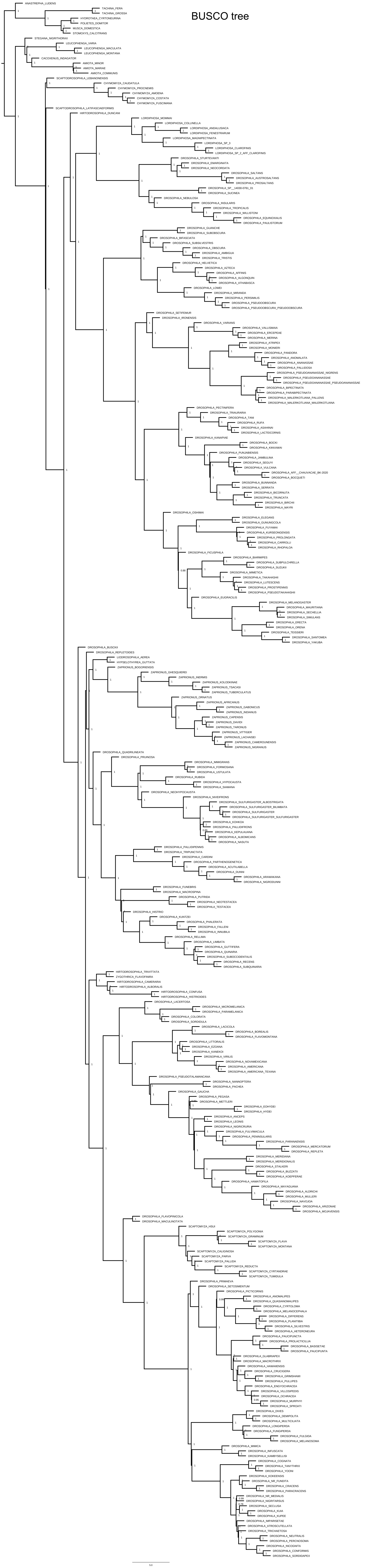

### S5 Fig

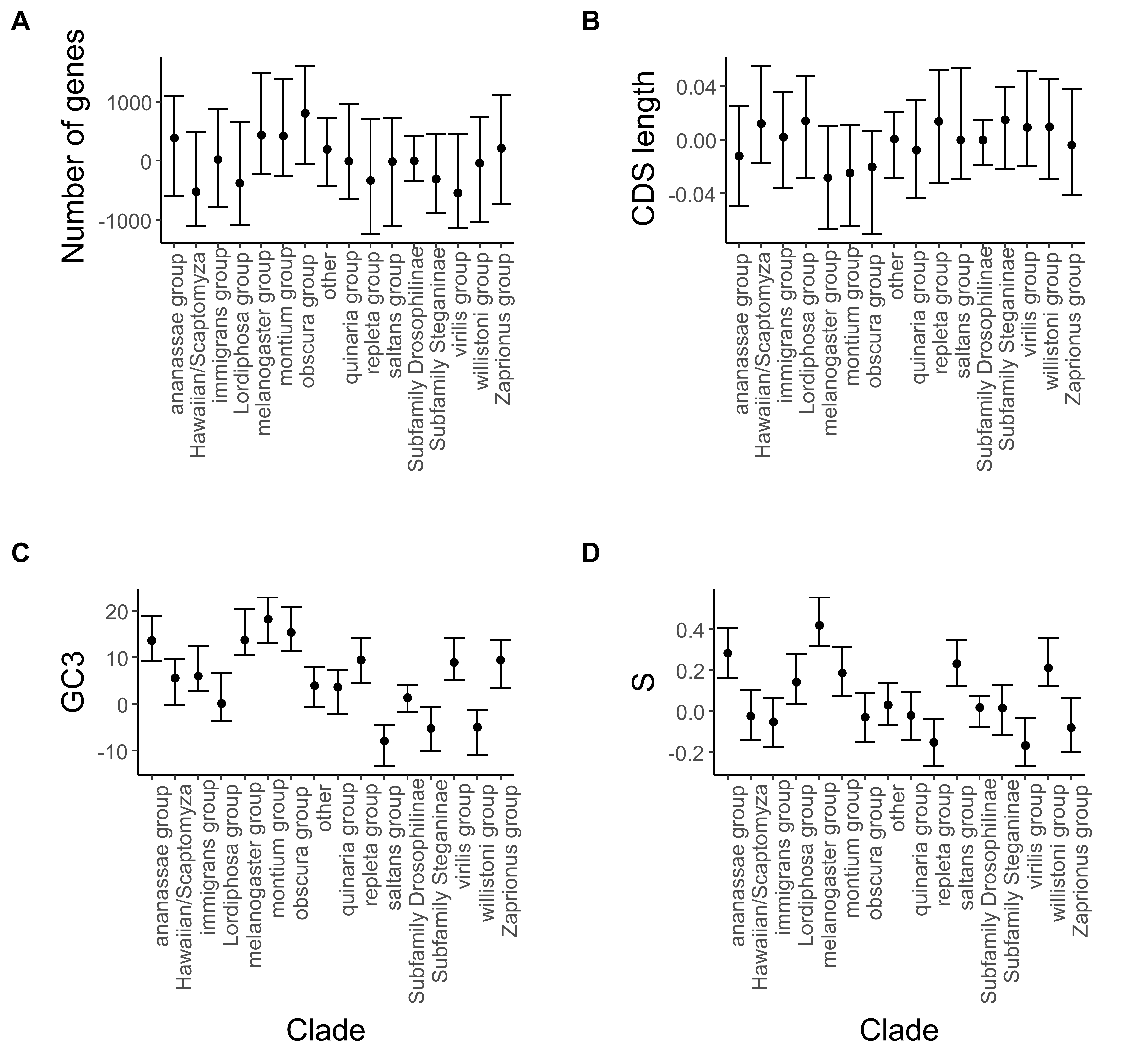
