## Supplementary material for "Comparative gene annotation and orthology assignments across 301 species of Drosophilidae": S1 File

### Content


- The model were specified as follows (MCMCglmm syntax)
- Model summaries (301 species)
    - Model for mean CDS length and gene number
    - Model for total CDS length and gene number
- Model summaries (215 species)
    - Model for mean CDS length and gene number
    - Model for total CDS length and gene number
- Model for CUB analysis (301 species)


### The model were specified as follows (MCMCglmm syntax):

```
# Model for mean CDS length and gene number
prior <- list(
  B = list(mu = rep(0, 14), V = diag(14) * 1e+10),  # Prior for fixed effects (mean 0, variance
  G = list(G1 = list(V=diag(2), nu=2, alpha.mu=rep(0,2), alpha.V=diag(2)*1000)),  # Prior for ra
  R = list(V=diag(2), nu=2.002)  # Prior for residual
)

# Fit the multivariate model
model <- MCMCglmm(cbind(mean_len, genes) ~ trait - 1 + trait:distances    + trait:RNA_seq + tra
       random = ~us(trait):Phylo,
                  rcov = ~us(trait):units,
                 family = rep("gaussian", 2),
                 ginverse = list(label = InverseTree),
                 prior = prior, data = annot_stat,
                 nitt = 1000000, burnin = 100000, thin = 1000, pr = TRUE)
```

Briefly, trait is a reserved variable that indexes columns of the response matrix in multi-response models, with -1 removing the global intercept so that each trait has its own baseline estimate. Fixed effects included phylogenetic distance from the reference species (trait:distances), availability of RNA-seq data (trait:RNA_seq), whether the species was itself a lift-over reference (trait:ref), assembled genome size (trait:genome_size), assembly contiguity (trait:ContigN50), and read-type (trait:Read_Types). The random effects term us(trait):Phylo describes the phylogenetic (co)variance matrix between gene length and gene number, and the residual variance term us(trait):units describes the residual covariance matrix. The argument ginverse fits a covariance structure among species to model non-independence due to common ancestry. Variance in CDS length and gene number were treated as Gaussian.

### Model summaries (301 species)

#### Model for mean CDS length and gene number

```
 Iterations = 100001:999001
 Thinning interval  = 1000
 Sample size  = 900 

 DIC: 4096.776 

 G-structure:  ~us(trait):label

                                  post.mean   l-95% CI   u-95% CI eff.samp
traitmean_len:traitmean_len.label  9.125e-04  1.522e-04  1.911e-03    900.0
traitgenes:traitmean_len.label    -1.996e+01 -4.005e+01 -3.771e+00    799.0
traitmean_len:traitgenes.label    -1.996e+01 -4.005e+01 -3.771e+00    799.0
traitgenes:traitgenes.label        5.069e+05  1.118e+05  9.474e+05    666.3

 R-structure:  ~us(trait):units

                                  post.mean   l-95% CI   u-95% CI eff.samp
traitmean_len:traitmean_len.units  8.231e-03  6.877e-03  9.527e-03     1098
traitgenes:traitmean_len.units    -1.764e+01 -2.668e+01 -8.321e+00      900
traitmean_len:traitgenes.units    -1.764e+01 -2.668e+01 -8.321e+00      900
traitgenes:traitgenes.units        7.145e+05  5.830e+05  8.657e+05      743

 Location effects: cbind(mean_len, genes) ~ trait - 1 + trait:distances + trait:RNA_seq + trait:ı

                              post.mean   l-95% CI   u-95% CI eff.samp   pMCMC   
traitmean_len                  1.690e+00  1.633e+00  1.756e+00    900.0 < 0.001 **
traitgenes                     1.266e+04  1.165e+04  1.382e+04    900.0 < 0.001 **
traitmean_len:distances        5.585e-02  2.378e-02  9.084e-02    900.0 0.00222 **
traitgenes:distances          -9.704e+02 -1.405e+03 -4.702e+02    900.0 < 0.001 **
traitmean_len:RNA_seqYes      -1.191e-02 -3.724e-02  1.710e-02    900.0 0.35556   
traitgenes:RNA_seqYes         -4.399e+02 -7.157e+02 -2.011e+02    900.0 0.00222 **
traitmean_len:refYes          -6.080e-03 -5.510e-02  4.016e-02    900.0 0.77333   
traitgenes:refYes              3.882e+01 -4.877e+02  4.942e+02    763.7 0.85556   
traitmean_len:genome_size     -6.613e-04 -9.138e-04 -4.026e-04    758.8 < 0.001 **
traitgenes:genome_size         1.345e+01  1.043e+01  1.651e+01    900.0 < 0.001 **
traitmean_len:ContigN50        1.108e-03 -4.511e-04  2.476e-03    900.0 0.11333   
traitgenes:ContigN50          -1.335e+01 -2.636e+01  5.950e-01    900.0 0.05111 . 
traitmean_len:Read_typesShort -5.995e-02 -1.068e-01 -2.026e-02    900.0 0.00444 **
```

```
traitgenes:Read_typesShort     1.024e+03  5.885e+02  1.459e+03    900.0 < 0.001 **
---
Signif. codes:  0 ‘***’ 0.001 ‘**’ 0.01 ‘*’ 0.05 ‘.’ 0.1 ‘ ’ 1

Mean and HPDinterval for phylogenetic heritability for mean_len:
[1] 0.09750539
          lower     upper
var1 0.01549091 0.1875801
attr(,"Probability")
[1] 0.95



Mean and HPDinterval for phylogenetic heritability for genes:
[1] 0.3991392
         lower     upper
var1 0.2053545 0.6380525
attr(,"Probability")
[1] 0.95
```

### Model for total CDS length and gene number

```
 Iterations = 100001:999001
 Thinning interval  = 1000
 Sample size  = 900

 DIC: 5237.656

 G-structure:  ~us(trait):label

                                    post.mean  l-95% CI  u-95% CI eff.samp
traittotal_cds:traittotal_cds.label 6.804e-01 1.693e-01 1.312e+00      900
traitgenes:traittotal_cds.label     4.846e+02 1.011e+02 9.857e+02      900
traittotal_cds:traitgenes.label     4.846e+02 1.011e+02 9.857e+02      900
traitgenes:traitgenes.label         5.325e+05 1.547e+05 9.946e+05      900

 R-structure:  ~us(trait):units

                                    post.mean  l-95% CI  u-95% CI eff.samp
traittotal_cds:traittotal_cds.units 1.097e+00 8.710e-01 1.306e+00    795.7
traitgenes:traittotal_cds.units     8.346e+02 6.829e+02 1.011e+03    810.4
traittotal_cds:traitgenes.units     8.346e+02 6.829e+02 1.011e+03    810.4
traitgenes:traitgenes.units         7.164e+05 5.799e+05 8.466e+05    900.0

 Location effects: cbind(total_cds, genes) ~ trait - 1 + trait:distances + trait:RNA_seq + trai

                                post.mean   l-95% CI   u-95% CI eff.samp   pMCMC
traittotal_cds                  2.147e+01  2.005e+01  2.264e+01      900 < 0.001 **
traitgenes                      1.275e+04  1.159e+04  1.379e+04      900 < 0.001 **
traittotal_cds:distances       -6.972e-01 -1.351e+00 -1.877e-01      900 0.02444 *
traitgenes:distances           -1.009e+03 -1.510e+03 -5.716e+02      900 < 0.001 **
traittotal_cds:RNA_seqYes      -7.677e-01 -1.094e+00 -4.726e-01     1044 < 0.001 **
traitgenes:RNA_seqYes          -4.455e+02 -7.178e+02 -2.015e+02     1074 < 0.001 **
traittotal_cds:refYes           6.206e-03 -5.651e-01  5.775e-01      900 0.98222
traitgenes:refYes               3.718e+01 -4.539e+02  4.629e+02      900 0.85111
traittotal_cds:genome_size      1.290e-02  9.038e-03  1.698e-02      900 < 0.001 **
traitgenes:genome_size          1.315e+01  9.935e+00  1.631e+01      900 < 0.001 **
traittotal_cds:ContigN50       -4.215e-03 -2.115e-02  1.514e-02      900 0.61333
traitgenes:ContigN50           -1.335e+01 -2.841e+01  5.558e-01      900 0.06667 .
traittotal_cds:Read_typesShort  6.924e-01  1.394e-01  1.202e+00      900 0.00889 **
traitgenes:Read_typesShort      1.059e+03  6.546e+02  1.507e+03      900 < 0.001 **
---
Signif. codes:  0 ‘***’ 0.001 ‘**’ 0.01 ‘*’ 0.05 ‘.’ 0.1 ‘ ’ 1
```

```
Mean and HPDinterval for phylogenetic heritability for total_cds:
[1] 0.3666086
         lower     upper
var1 0.1642473 0.6026227
attr(,"Probability")
[1] 0.95


Mean and HPDinterval for phylogenetic heritability for genes:
[1] 0.4100006
         lower     upper
var1 0.2046233 0.6158211
attr(,"Probability")
[1] 0.95
```

### Model summaries (215 species)

#### Model for mean CDS length and gene number

```
 Iterations = 100001:999001
 Thinning interval  = 1000
 Sample size  = 900 

 DIC: 2717.105 

 G-structure:  ~us(trait):label

                                   post.mean   l-95% CI   u-95% CI eff.samp
traitmean_len:traitmean_len.label  7.942e-04  2.971e-05  1.837e-03      900
traitgenes:traitmean_len.label    -1.468e+01 -2.772e+01 -2.955e+00      900
traitmean_len:traitgenes.label    -1.468e+01 -2.772e+01 -2.955e+00      900
traitgenes:traitgenes.label        3.410e+05  1.743e+05  5.434e+05     1061

 R-structure:  ~us(trait):units

                                   post.mean   l-95% CI   u-95% CI eff.samp
traitmean_len:traitmean_len.units  1.050e-02  8.396e-03  1.240e-02    900.0
traitgenes:traitmean_len.units    -6.930e+00 -1.296e+01 -4.399e-02    729.7
traitmean_len:traitgenes.units    -6.930e+00 -1.296e+01 -4.399e-02    729.7
traitgenes:traitgenes.units        2.009e+05  1.532e+05  2.479e+05    900.0

 Location effects: cbind(mean_len, genes) ~ trait - 1 + trait:distances + trait:RNA_seq + trait

                           post.mean   l-95% CI   u-95% CI eff.samp   pMCMC   
traitmean_len              1.686e+00  1.599e+00  1.770e+00    900.0 < 0.001 **
traitgenes                 1.263e+04  1.180e+04  1.348e+04    900.0 < 0.001 **
traitmean_len:distances    6.692e-02  2.558e-02  1.107e-01    968.2 0.00444 **
traitgenes:distances      -9.152e+02 -1.329e+03 -5.442e+02    900.0 < 0.001 **
traitmean_len:RNA_seqYes  -1.593e-02 -4.935e-02  1.658e-02   1006.2 0.39556   
traitgenes:RNA_seqYes     -1.411e+02 -3.064e+02  1.898e+01    900.0 0.09556 . 
traitmean_len:refYes       1.375e-03 -5.012e-02  5.909e-02   1067.0 0.99778   
traitgenes:refYes         -1.315e+02 -3.999e+02  1.314e+02    900.0 0.34889   
traitmean_len:genome_size -5.945e-04 -9.873e-04 -1.971e-04    779.0 0.00444 **
traitgenes:genome_size     1.249e+01  1.016e+01  1.489e+01    800.8 < 0.001 **
traitmean_len:ContigN50    5.192e-04 -1.271e-03  2.389e-03    900.0 0.56667   
traitgenes:ContigN50      -8.653e+00 -1.817e+01  6.128e-01    679.8 0.07556 . 
---
```

```
Signif. codes:  0 ‘***’ 0.001 ‘**’ 0.01 ‘*’ 0.05 ‘.’ 0.1 ‘ ’ 1


Mean and HPDinterval for phylogenetic heritability for mean_len:
[1] 0.06884757
           lower     upper
var1 0.002767685 0.1524913
attr(,"Probability")
[1] 0.95

Mean and HPDinterval for phylogenetic heritability for genes:
[1] 0.6174786
         lower     upper
var1 0.4747389 0.7780925
attr(,"Probability")
[1] 0.95
```

### Model for total CDS length and gene number

```
 Iterations = 100001:999001
 Thinning interval  = 1000
 Sample size  = 900

 DIC: 3316.107

 G-structure:  ~us(trait):label

                                    post.mean  l-95% CI  u-95% CI eff.samp
traittotal_cds:traittotal_cds.label 4.508e-01 2.125e-01 7.390e-01    991.6
traitgenes:traittotal_cds.label     3.407e+02 1.427e+02 5.570e+02   1023.6
traittotal_cds:traitgenes.label     3.407e+02 1.427e+02 5.570e+02   1023.6
traitgenes:traitgenes.label         3.360e+05 1.676e+05 5.501e+05   1004.2

 R-structure:  ~us(trait):units

                                    post.mean  l-95% CI  u-95% CI eff.samp
traittotal_cds:traittotal_cds.units 2.743e-01 2.112e-01 3.393e-01     1135
traitgenes:traittotal_cds.units     2.070e+02 1.626e+02 2.661e+02     1104
traittotal_cds:traitgenes.units     2.070e+02 1.626e+02 2.661e+02     1104
traitgenes:traitgenes.units         2.027e+05 1.578e+05 2.513e+05      900

 Location effects: cbind(total_cds, genes) ~ trait - 1 + trait:distances + trait:RNA_seq + trai

                            post.mean   l-95% CI   u-95% CI eff.samp  pMCMC
traittotal_cds              2.146e+01  2.043e+01  2.246e+01    900.0 <0.001 **
traitgenes                  1.263e+04  1.176e+04  1.349e+04   1085.9 <0.001 **
traittotal_cds:distances   -5.651e-01 -1.029e+00 -1.374e-01   1034.5 0.0178 *
traitgenes:distances       -9.111e+02 -1.288e+03 -5.537e+02   1020.8 <0.001 **
traittotal_cds:RNA_seqYes  -3.751e-01 -5.610e-01 -1.815e-01    900.0 <0.001 **
traitgenes:RNA_seqYes      -1.429e+02 -3.296e+02  1.518e+01    972.8 0.0867 .
traittotal_cds:refYes      -1.973e-01 -5.241e-01  1.146e-01    900.0 0.2644
traitgenes:refYes          -1.313e+02 -4.229e+02  1.304e+02    900.0 0.3733
traittotal_cds:genome_size  1.263e-02  9.622e-03  1.537e-02    900.0 <0.001 **
traitgenes:genome_size      1.247e+01  9.944e+00  1.496e+01    900.0 <0.001 **
traittotal_cds:ContigN50   -4.576e-03 -1.575e-02  6.500e-03    900.0 0.4689
traitgenes:ContigN50       -8.867e+00 -1.963e+01 -4.223e-01    900.0 0.0667 .
---
Signif. codes:  0 ‘***’ 0.001 ‘**’ 0.01 ‘*’ 0.05 ‘.’ 0.1 ‘ ’ 1
```

```
Mean and HPDinterval for phylogenetic heritability for total_cds:
[1] 0.609703
         lower    upper
var1 0.4540152 0.768885
attr(,"Probability")
[1] 0.95


Mean and HPDinterval for phylogenetic heritability for genes:
[1] 0.6118986
        lower     upper
var1 0.451971 0.7716967
attr(,"Probability")
[1] 0.95
```

### Model for CUB analysis (301 species)

```
prior <- list(R = list(V = diag(7), nu = 0.002),
              G = list(G1 = list(V = diag(7), nu = 0.002)))

# Fit the MCMCglmm model
model <- MCMCglmm(cbind(mean_gc3, GC_nonCoding, S, genome_size, PC1_AA, PC2_AA, NC) ~ trait - 1,
                  random = ~us(trait):label,
                  rcov = ~us(trait):units,
                  family = rep("gaussian", 7),
                  ginverse = list(label = InverseTree),
                  data = cub_df,
                  prior = prior,
                  nitt = 100000,
                  burnin = 10000,
                  thin = 100,
                  pr=TRUE)

> summary(model)

 Iterations = 10001:99901
 Thinning interval  = 100
 Sample size  = 900

 DIC: -4341.228

 G-structure:  ~us(trait):label

                                           post.mean   l-95% CI   u-95% CI eff.samp
traitmean_gc3:traitmean_gc3.label          5.062e-01  4.292e-01  5.925e-01    900.0
traitGC_nonCoding:traitmean_gc3.label      1.588e-01  1.181e-01  1.983e-01    900.0
traitS:traitmean_gc3.label                -2.649e-03 -4.026e-03 -9.763e-04    900.0
traitgenome_size:traitmean_gc3.label      -4.850e-01 -1.304e+00  2.880e-01   1031.4
traitPC1_AA:traitmean_gc3.label           -7.822e-02 -9.392e-02 -6.263e-02    900.0
traitPC2_AA:traitmean_gc3.label            1.053e-01  8.191e-02  1.283e-01    900.0
traitNC:traitmean_gc3.label                3.226e-05 -2.402e-04  3.413e-04    900.0
traitmean_gc3:traitGC_nonCoding.label      1.588e-01  1.181e-01  1.983e-01    900.0
traitGC_nonCoding:traitGC_nonCoding.label  2.008e-01  1.687e-01  2.376e-01    593.9
traitS:traitGC_nonCoding.label            -2.641e-03 -3.758e-03 -1.596e-03    984.3
traitgenome_size:traitGC_nonCoding.label   8.779e-02 -5.651e-01  7.561e-01    565.6
traitPC1_AA:traitGC_nonCoding.label       -3.337e-02 -4.349e-02 -2.374e-02    582.0
traitPC2_AA:traitGC_nonCoding.label        2.151e-02  7.765e-03  3.674e-02    708.7
```

```
traitNC:traitGC_nonCoding.label             1.422e-05 -1.769e-04  1.936e-04    900.0
traitmean_gc3:traitS.label                 -2.649e-03 -4.026e-03 -9.763e-04    900.0
traitGC_nonCoding:traitS.label             -2.641e-03 -3.758e-03 -1.596e-03    984.3
traitS:traitS.label                         3.240e-04  2.657e-04  3.848e-04    900.0
traitgenome_size:traitS.label               2.726e-02  3.175e-03  5.202e-02    900.0
traitPC1_AA:traitS.label                    1.856e-04 -2.215e-04  5.558e-04    990.5
traitPC2_AA:traitS.label                   -6.547e-04 -1.215e-03 -8.387e-05    900.0
traitNC:traitS.label                       -4.758e-07 -7.644e-06  7.726e-06    900.0
traitmean_gc3:traitgenome_size.label       -4.850e-01 -1.304e+00  2.880e-01   1031.4
traitGC_nonCoding:traitgenome_size.label    8.779e-02 -5.651e-01  7.561e-01    565.6
traitS:traitgenome_size.label               2.726e-02  3.175e-03  5.202e-02    900.0
traitgenome_size:traitgenome_size.label     7.428e+01  5.553e+01  9.753e+01    900.0
traitPC1_AA:traitgenome_size.label         -5.403e-02 -2.445e-01  1.627e-01    658.5
traitPC2_AA:traitgenome_size.label         -1.608e-01 -4.683e-01  1.561e-01    900.0
traitNC:traitgenome_size.label             -1.261e-04 -3.552e-03  3.889e-03   1038.4
traitmean_gc3:traitPC1_AA.label            -7.822e-02 -9.392e-02 -6.263e-02    900.0
traitGC_nonCoding:traitPC1_AA.label        -3.337e-02 -4.349e-02 -2.374e-02    582.0
traitS:traitPC1_AA.label                    1.856e-04 -2.215e-04  5.558e-04    990.5
traitgenome_size:traitPC1_AA.label         -5.403e-02 -2.445e-01  1.627e-01    658.5
traitPC1_AA:traitPC1_AA.label               2.878e-02  2.461e-02  3.389e-02   1102.8
traitPC2_AA:traitPC1_AA.label              -7.399e-03 -1.274e-02 -2.968e-03    990.2
traitNC:traitPC1_AA.label                  -2.277e-06 -7.951e-05  6.820e-05    900.0
traitmean_gc3:traitPC2_AA.label             1.053e-01  8.191e-02  1.283e-01    900.0
traitGC_nonCoding:traitPC2_AA.label         2.151e-02  7.765e-03  3.674e-02    708.7
traitS:traitPC2_AA.label                   -6.547e-04 -1.215e-03 -8.387e-05    900.0
traitgenome_size:traitPC2_AA.label         -1.608e-01 -4.683e-01  1.561e-01    900.0
traitPC1_AA:traitPC2_AA.label              -7.399e-03 -1.274e-02 -2.968e-03    990.2
traitPC2_AA:traitPC2_AA.label               5.670e-02  4.806e-02  6.869e-02   1057.0
traitNC:traitPC2_AA.label                   1.038e-05 -9.595e-05  1.058e-04    704.7
traitmean_gc3:traitNC.label                 3.226e-05 -2.402e-04  3.413e-04    900.0
traitGC_nonCoding:traitNC.label             1.422e-05 -1.769e-04  1.936e-04    900.0
traitS:traitNC.label                       -4.758e-07 -7.644e-06  7.726e-06    900.0
traitgenome_size:traitNC.label             -1.261e-04 -3.552e-03  3.889e-03   1038.4
traitPC1_AA:traitNC.label                  -2.277e-06 -7.951e-05  6.820e-05    900.0
traitPC2_AA:traitNC.label                   1.038e-05 -9.595e-05  1.058e-04    704.7
traitNC:traitNC.label                       1.054e-05  8.625e-06  1.241e-05    706.7

 R-structure:  ~us(trait):units

                                        post.mean   l-95% CI   u-95% CI eff.samp
traitmean_gc3:traitmean_gc3.units       1.168e-02  1.108e-03 2.904e-02    72.37
traitGC_nonCoding:traitmean_gc3.units  -2.752e-03 -2.109e-02 1.627e-02   178.93
traitS:traitmean_gc3.units             -2.169e-04 -1.106e-03 5.685e-04    87.11
```

| | | | | |
|---|---|---|---|---|
| traitgenome_size:traitmean_gc3.units | 1.176e-01 | -1.799e+00 | 2.321e+00 | 63.14 |
| traitPC1_AA:traitmean_gc3.units | -1.248e-04 | -5.734e-03 | 5.414e-03 | 167.41 |
| traitPC2_AA:traitmean_gc3.units | -9.076e-03 | -2.772e-02 | 1.638e-02 | 119.53 |
| traitNC:traitmean_gc3.units | 7.824e-07 | -8.198e-05 | 9.433e-05 | 1064.55 |
| traitmean_gc3:traitGC_nonCoding.units | -2.752e-03 | -2.109e-02 | 1.627e-02 | 178.93 |
| traitGC_nonCoding:traitGC_nonCoding.units | 3.860e-02 | 9.667e-03 | 8.100e-02 | 168.87 |
| traitS:traitGC_nonCoding.units | 5.913e-04 | -4.895e-04 | 1.568e-03 | 516.54 |
| traitgenome_size:traitGC_nonCoding.units | -1.572e-01 | -2.022e+00 | 1.379e+00 | 144.67 |
| traitPC1_AA:traitGC_nonCoding.units | -7.596e-03 | -1.652e-02 | 1.193e-03 | 192.66 |
| traitPC2_AA:traitGC_nonCoding.units | 3.004e-02 | 8.778e-03 | 5.225e-02 | 273.90 |
| traitNC:traitGC_nonCoding.units | -2.944e-06 | -1.752e-04 | 1.589e-04 | 900.00 |
| traitmean_gc3:traitS.units | -2.169e-04 | -1.106e-03 | 5.685e-04 | 87.11 |
| traitGC_nonCoding:traitS.units | 5.913e-04 | -4.895e-04 | 1.568e-03 | 516.54 |
| traitS:traitS.units | 2.082e-04 | 1.244e-04 | 2.985e-04 | 900.00 |
| traitgenome_size:traitS.units | -6.790e-02 | -1.418e-01 | 1.125e-02 | 760.48 |
| traitPC1_AA:traitS.units | -5.797e-05 | -3.527e-04 | 3.574e-04 | 677.53 |
| traitPC2_AA:traitS.units | 1.010e-03 | 2.699e-05 | 2.056e-03 | 900.00 |
| traitNC:traitS.units | -1.735e-08 | -1.221e-05 | 1.244e-05 | 900.00 |
| traitmean_gc3:traitgenome_size.units | 1.176e-01 | -1.799e+00 | 2.321e+00 | 63.14 |
| traitGC_nonCoding:traitgenome_size.units | -1.572e-01 | -2.022e+00 | 1.379e+00 | 144.67 |
| traitS:traitgenome_size.units | -6.790e-02 | -1.418e-01 | 1.125e-02 | 760.48 |
| traitgenome_size:traitgenome_size.units | 2.706e+02 | 1.832e+02 | 3.616e+02 | 900.00 |
| traitPC1_AA:traitgenome_size.units | -8.617e-02 | -6.370e-01 | 4.609e-01 | 168.76 |
| traitPC2_AA:traitgenome_size.units | -9.281e-01 | -2.069e+00 | 5.233e-02 | 253.22 |
| traitNC:traitgenome_size.units | 4.402e-05 | -1.408e-02 | 1.504e-02 | 900.00 |
| traitmean_gc3:traitPC1_AA.units | -1.248e-04 | -5.734e-03 | 5.414e-03 | 167.41 |
| traitGC_nonCoding:traitPC1_AA.units | -7.596e-03 | -1.652e-02 | 1.193e-03 | 192.66 |
| traitS:traitPC1_AA.units | -5.797e-05 | -3.527e-04 | 3.574e-04 | 677.53 |
| traitgenome_size:traitPC1_AA.units | -8.617e-02 | -6.370e-01 | 4.609e-01 | 168.76 |
| traitPC1_AA:traitPC1_AA.units | 3.981e-03 | 1.082e-03 | 7.363e-03 | 336.49 |
| traitPC2_AA:traitPC1_AA.units | -4.965e-03 | -1.240e-02 | 1.957e-03 | 276.77 |
| traitNC:traitPC1_AA.units | -2.991e-08 | -6.387e-05 | 5.107e-05 | 900.00 |
| traitmean_gc3:traitPC2_AA.units | -9.076e-03 | -2.772e-02 | 1.638e-02 | 119.53 |
| traitGC_nonCoding:traitPC2_AA.units | 3.004e-02 | 8.778e-03 | 5.225e-02 | 273.90 |
| traitS:traitPC2_AA.units | 1.010e-03 | 2.699e-05 | 2.056e-03 | 900.00 |
| traitgenome_size:traitPC2_AA.units | -9.281e-01 | -2.069e+00 | 5.233e-02 | 253.22 |
| traitPC1_AA:traitPC2_AA.units | -4.965e-03 | -1.240e-02 | 1.957e-03 | 276.77 |
| traitPC2_AA:traitPC2_AA.units | 6.391e-02 | 4.222e-02 | 8.697e-02 | 511.66 |
| traitNC:traitPC2_AA.units | 1.212e-05 | -1.976e-04 | 2.158e-04 | 900.00 |
| traitmean_gc3:traitNC.units | 7.824e-07 | -8.198e-05 | 9.433e-05 | 1064.55 |
| traitGC_nonCoding:traitNC.units | -2.944e-06 | -1.752e-04 | 1.589e-04 | 900.00 |
| traitS:traitNC.units | -1.735e-08 | -1.221e-05 | 1.244e-05 | 900.00 |
| traitgenome_size:traitNC.units | 4.402e-05 | -1.408e-02 | 1.504e-02 | 900.00 |

```
traitPC1_AA:traitNC.units                    -2.991e-08 -6.387e-05 5.107e-05   900.00
traitPC2_AA:traitNC.units                     1.212e-05 -1.976e-04 2.158e-04   900.00
traitNC:traitNC.units                         2.040e-05  1.637e-05 2.582e-05   900.00

 Location effects: cbind(mean_gc3, GC_nonCoding, S, genome_size, PC1_AA, PC2_AA, NC) ~ trait - :

                  post.mean l-95% CI u-95% CI eff.samp  pMCMC
traitmean_gc3       51.4505  47.3798  56.1655      900 <0.001 **
traitGC_nonCoding   33.4987  30.7485  36.3058      900 <0.001 **
traitS               0.4607   0.3523   0.5718      900 <0.001 **
traitgenome_size   228.5554 178.2836 280.6299      900 <0.001 **
traitPC1_AA          1.6399   0.5880   2.6138      900 <0.001 **
traitPC2_AA         -2.4431  -3.8626  -1.0000      900 <0.001 **
traitNC              0.3011   0.2823   0.3209      900 <0.001 **
---
Signif. codes:  0 '***' 0.001 '**' 0.01 '*' 0.05 '.' 0.1 ' ' 1

> mean(heritability_mean_gc3)
[1] 0.9995968
> HPDinterval(heritability_mean_gc3)
         lower     upper
var1 0.9990681 0.9999826
attr(,"Probability")
[1] 0.95
> mean(heritability_S)
[1] 0.98878
> HPDinterval(heritability_S)
         lower    upper
var1 0.9827214 0.994018
attr(,"Probability")
[1] 0.95
```
